# ABCC6 and ANK regulate extracellular homeostasis of pyrophosphate and citrate and affect mineral deposition in bones and soft connective tissues

**DOI:** 10.1101/2025.10.09.681172

**Authors:** Flora Szeri, Fatemeh Niaziorimi, Celina Ng, Cassandra Mikkelson, Ibtesam Rajpar, Qinglin Wu, Steven P. Matyus, Margery A. Connelly, George Tavadze, Stefan Lundkvist, Ryan E. Tomlinson, Koen van de Wetering

## Abstract

ABCC6 and ANK are integral membrane proteins involved in the efflux of specific organic anions. ABCC6, primarily expressed in the liver, regulates circulating levels of the mineralization inhibitor pyrophosphate (PPi). In contrast, ANK is widely expressed and plays a dual role, maintaining extracellular PPi homeostasis and, notably, mediating cellular citrate efflux. We studied how both proteins affected extracellular metabolite levels, ectopic calcification, and bone homeostasis.

We observed that plasma PPi was reduced by ∼36% in *Ank^ank/ank^*mice and ∼60% in *Abcc6^−/−^* mice. However, plasma citrate levels depended primarily on ANK, dropping ∼75% in *Ank^ank/ank^* and double mutants, but remaining unchanged in *Abcc6^−/−^* mice.

MicroCT revealed extreme ectopic calcification in double mutants, far exceeding either single knockout, affecting muzzle skin and ear cartilage. Oral citrate was bioavailable and, at high doses, prevented soft tissue calcification in *Abcc6^−/−^* mice, suggesting a systemic protective role.

In bone, ANK was essential for incorporating both PPi and citrate, while ABCC6 mainly affected PPi. ANK deficiency led to reduced trabecular volume, cortical thickness, cortical area fraction, and mineral density, with more pronounced effects in males. Biomechanical testing showed decreased ultimate moment, bending rigidity, and energy in ANK-deficient femora, alongside increased post-yield displacement, indicating compensatory matrix changes.

Collectively, our findings identify ANK as a dual regulator of PPi and citrate, with a previously unrecognized role in preventing soft tissue calcification. This study positions ANK as a potential therapeutic target for mineralization disorders like pseudoxanthoma elasticum (caused by ABCC6 deficiency) and conditions of low bone mineral density like osteoporosis.

## Introduction

Physiological mineralization is regulated by the coordinated action of factors that promote and inhibit calcium phosphate precipitation. Under normal conditions, mineralization, mainly in the form of calcium phosphate crystals, is restricted to specific sites of the body, including bones and teeth^1–4^. Calcium and phosphate are present at high concentrations in the extracellular environment allowing the coordinated, physiological deposition of calcium phosphate in the collagen-rich osteoid of developing bone tissue. Intriguingly, calcium and phosphate are also present at high concentrations in tissues that under normal conditions do not mineralize. These soft tissues are protected against ectopic deposition of calcium phosphate crystals by complementary, inhibitory factors^5^.

Inorganic pyrophosphate (PPi) is a crucial mineralization regulator and its absence results in widespread calcification of soft connective tissues^6–8^. Extracellular levels of PPi (ePPi) are tightly controlled to on the one hand allow mineralization of bones and teeth, while on the other hand preventing the ectopic calcification of soft connective tissues. All ePPi depends on the activity of the ectoenzyme ectonucleotide pyrophosphatase phosphodiesterase 1 (ENPP1), which converts ATP and other nucleoside triphosphates (NTPs) into PPi and their respective nucleoside monophosphate (NMP)^7^. We have recently identified two proteins important for the release of ATP from cells for ENPP1-mediated PPi generation in the extracellular environment. In 2014 we found that ATP Binding Cassette Subfamily C Member 6 (ABCC6), which is predominantly expressed in the liver, mediates release of ATP from hepatocytes into the blood circulation, a process underlying 60-70% of all PPi present in plasma^9^. More recently, we identified Progressive Ankylosis Protein (ANK) as another crucial pathway for cellular ATP release and extracellular PPi formation^10,11^. ANK is ubiquitous and found at especially high levels in osteoblasts and osteocytes, where it is responsible for incorporation of PPi into the mineral phase of bone^11–13^. ANK-mediated ATP release is also crucial for ePPi formation in tissues and body fluids not in direct contact with the blood circulation, including the intervertebral disc, cartilage, and synovial fluid^14,15^. In addition, ANK activity also underlies about 20% of the ePPi present in plasma^11^. Notably, ANK also mediates the release of several TCA cycle intermediates, of which citrate is quantitatively the most important^10^. This is important as citrate chelates calcium and might contribute to the mineralization inhibitory effects of ANK.

A complex, incompletely characterized, network of proteins governs ePPi homeostasis^7^. Other proteins participating in ePPi homeostasis are tissue non-specific alkaline phosphatase (TNAP), which degrades PPi into organic phosphate (Pi) and CD73. The latter converts adenosine monophosphate (AMP) into Pi and the TNAP inhibitor adenosine^7^.

Biallelic mutations in the genes encoding proteins involved in ePPi homeostasis are associated with specific mineralization disorders. Absence of ENPP1 results in generalized calcification of infancy (GACI)^16,17^, a condition that becomes life-threatening shortly after birth because of extensive calcification of major arteries. Plasma of GACI patients is virtually devoid of PPi, explaining the severity of the disease. Pseudoxanthoma elasticum (PXE) is a late onset, slowly progressive mineralization disorder caused by mutations in ABCC6^18^. Ectopic mineral deposits in PXE patients are noted in skin, eyes, and arteries and the first clinical manifestations are most often encountered in the second decade of life. Patients with inactivating mutations in ANK suffer from ankylosis of their spine and calcium phosphate deposits in joints of hand and feet^15,19^. Insufficient TNAP activity results in a condition known as hypophosphatasia in which accumulation of ePPi results in hypo-mineralization of bones^20,21^. In patients with mutations in the *NT5E* gene, which encodes CD73, reduced conversion of AMP into adenosine results in increased TNAP activity, low plasma ePPi concentrations and, consequently, ectopic calcification of major arteries and the capsule of joints^22^.

In the current study, we first delineated how ANK and ABCC6 affect ePPi concentrations in plasma and the mineral phase bone using the recently generated *Abcc6^−/−^*;*Ank^ank/ank^* double mutant mice. Next, we determined how both proteins on the one hand contribute to prevention of ectopic calcification of soft tissues while on the other hand regulating bone structural and mechanical properties.

## Materials and methods

### Reagents

Unless otherwise indicated all reagents were obtained from Fisher Scientific (Waltham, MA).

#### Animals

*Abcc6^−/−^*;*Ank^ank/ank^* double mutant mice were generated by crossing *Abcc6^−/−^* mice^23^ with mice harboring the progressive ankylosis allele (*ank*, obtained from The Jackson Laboratory (Bar Harbor, ME; C3FeB6 *A/A^w-J^-Ank^ank/J^*, stock number 000200). All strains were crossed for >14 generations with C57Bl/6J animals. This is important as we found that a pathogenic mutation at position c.1863 in the Abcc6 allele was present in the C3FeB6 *A/A^w-J^-Ank^ank/J^*animals imported from the Jackson Laboratory (G->A). Animals used in our studies came from *Abcc6^+/+^*;*Ank^ank/wt^* and *Abcc6^−/−^*;*Ank^ank/wt^*breeding pairs.

In pharmacokinetic studies, 300 mM tri-potassium citrate was administered by oral gavage at a dose of 10 µl/g body weight. At the indicated times, blood was collected by cardiac puncture in syringes containing 25 µl of an 8.7% EDTA solution. Plasma was subsequently prepared by centrifugation for 10 minutes at 6000 RCF and 4 °C. In experiments designed to determine the effect on ectopic calcification, citrate treatment started directly following weaning, when *Abcc6^−/−^* mice are still free of muzzle skin calcification. Tri-potassium citrate was provided continuously via the drinking water at concentrations of 40 or 80 mM. Animals in the control cohort received regular water. Treatment was continued for 6 months, and water was replenished twice per week. At the end of the experiment animals were sacrificed and tissues collected to quantify ectopic calcification by micro-computed tomography (microCT).

Animal studies were approved by the Institutional Animal Care and Use Committee of Thomas Jefferson University in accordance with the National Institutes of Health Guide for Care and Use of Laboratory Animals under approval numbers 02081-2 and 02274-2.

#### Quantification of PPi in plasma and bone

For plasma PPi analysis blood was collected from 3-month-old animals. Approximately 800 µl of blood was collected by cardiac puncture in syringes containing 100 µl of a CTAD/EDTA (75 µl CTAD (Greiner Bio-One, Vacuette Tube, product # 454064) and 25 µl of an 8.7% EDTA solution) mixture. Plasma was prepared by centrifugation (10 min, 4 °C, 1500 RCF). Platelets were subsequently removed from plasma by filtration over a Centrisart I (300,000 mwco) filtration device, as described previously^24^. Plasma was stored at −80 °C until analysis.

Bones of 3-month-old animals were processed for PPi analysis as previously described^11^, with modifications. In short, 1000 µl 10% formic acid was added per 25 mg of bone from which bone marrow was removed by centrifugation. Samples we subsequently incubated overnight at 55 °C. After centrifugation (30,000 RCF, 4 °C, 10 min), the supernatant was stored at −80 °C until analysis. Bone extracts were diluted 100-fold in water before analysis.

PPi was quantified in plasma and diluted bone extracts as described previously^9,10^, with small modifications. 2.5 µl of sample was incubated in a total volume of 80 µl of reaction mixture containing 75 mU/mL ATP sulfurylase (ATPS, New England Biolabs, Cambridge, MA, USA), 1 μmol/L adenosine phosphosulfate (APS, Santa Cruz Biotechnology), 80 μmol/L MgCl_2_, and 50 mmol/L HEPES (pH 7.4). After incubation for 30 minutes at 30 °C and 10 minutes at 90 °C, 10 μL of the reaction mixture was incubated with 30 μL of SL reagent (BioThema, Handen, Sweden). Luminescence was quantified in a Spark Tecan plate reader (Tecan, Männedorf, Switzerland). PPi concentrations were finally calculated by taking along a PPi standard curve. Studies included similar numbers of male and female mice for each genotype.

### Quantification of citrate in plasma and bone

Blood was collected in a syringe containing 25 µl of an 8.6 % tri-potassium EDTA solution and stored on ice till further processing. Plasma was prepared by centrifugation (10 minutes, 4 °C, 6000 RCF) and stored at −80 °C till analysis.

Citrate was quantified in tibiae from which bone marrow was removed by centrifugation. Bone was subsequently dissolved in 10% formic acid (1 ml/25 mg bone) and stored at −80 °C till analyzed by Labcorp (Morrisville, NC, USA) for testing on a Vantera^®^ Clinical Analyzer. Citrate levels were determined using NMR spectroscopy as previously described ^25,26^. Inter-assay imprecision studies revealed coefficients of variation (%CV) for citrate range from 5.2% to 9.6% for high and low concentration samples, respectively. For the bone extracted specimens, 40 µL was aspirated and diluted 8-fold with NMR diluent 1 ^25^ onboard the Vantera^®^ Clinical Analyzer before acquisition of the NMR spectra. NMR spectra were transferred to and the data were processed by an in-house developed MATLAB® program (Mathworks, Natick, MA). Given the acidic pH of the samples, the location of the citrate signal peaks in the NMR spectrum was significantly different between bone extracts and EDTA plasma. Therefore, a restricted region around each citrate peak was selected for citrate quantification using a non-negative linear least squares deconvolution algorithm. Conversion factors were used to calculate the citrate concentration by multiplying the sum of the coefficients derived from the deconvolution. The conversion factors were validated from standard curves carried out under similar conditions with known citrate concentrations.

### MicroCT

Each bone was scanned using a Bruker Skyscan 1275 microCT system (Bruker, Billerica, MA) equipped with a 1 mm aluminum filter. One femur from each mouse was scanned at 55 kV and 181 µA with a 74 ms exposure time. Transverse scan slices were obtained by placing the long axis of the bone parallel to the z axis of the scanner using a custom 3D printed sample holder. An isometric voxel size of 13 µm was used. Images were reconstructed using nRecon (Bruker, Billerica, MA) and analyzed using CTan (Bruker, Billerica, MA). Cortical bone was analyzed using a 1 mm thick region of interest centered at the mid-diaphysis of the femur. Quantitative analysis was performed in accordance with the recommendations of the American Society for Bone and Mineral Research^27^.

Ectopic calcification in the muzzle skin and ears was determined by µCT at 10 µm resolution using the same settings as given above for bone. Mineral deposits were subsequently quantified in reconstructed images using nRecon and CTan software (Bruker, Billerica, MA) using a lower cut-off value of 55 (BMD of 0.17 g/cm3)

### Three-point Bending assay

Three-point bending was performed on bones that had been stored at −20 °C in PBS-soaked gauze after harvest, as described previously^10^. Femora were scanned with microCT before performing three-point bending. Briefly, each femur was oriented on a standard fixture with femoral condyles facing down and a bending span of 8.7 mm. Next, a monotonic displacement ramp of 0.1 mm/s was applied until failure, with force and displacement acquired digitally. The force-displacement curves were converted to stress-strain using microCT-based geometry and analyzed using a custom GNU Octave script.

### Statistical Analysis

All the statistical analyses were performed using Prism version 10.6.1 (GraphPad Software, San Diego, CA, USA) and are presented as individual values and mean ± SD or as percentages ± SD. Statistical analysis was performed by Brown-Forsythe and Welch’s ANOVA followed by Dunnett’s multiple comparison test.

## Results

### Plasma pyrophosphate levels depend on ABCC6 and ANK

Previous work from our laboratory indicated that ABCC6 and ANK underlie 60 and 20%, respectively, of the amounts of PPi present in plasma^10,11^. In our current study levels of PPi in *Ank^ank/ank^* mice were 36% lower than in plasma of wild type control mice (mean 1.2 ± 0.5 µM versus 1.9 ± 0.6µM in wild type plasma) (Fig. 1A). As expected, PPi concentrations in plasma of *Abcc6^−/−^* mice were much lower (0.7 ± 0.2 µM), ∼ 40% of wild type levels. These values were in line with previously reported data from our group^10,11^. As both ABCC6 and ANK contribute to plasma PPi, we anticipated even lower concentrations of PPi in *Abcc6^−/−^*;*Ank^ank/ank^*mice. Unexpectedly, PPi plasma concentrations did not substantially differ between *Abcc6^−/−^*;*Ank^ank/ank^* and *Abcc6^−/−^* mice (mean 0.6 ± 0.3 µM versus 0.7 ± 0.2 µM, in *Abcc6^−/−^* mice), however. PPi plasma concentrations can be affected by small differences in blood collection and plasma preparation. For instance, platelets release ATP upon activation, which can be converted into PPi by the soluble ENPP1 that is present in plasma. Platelet factor-4 (PF-4) is an indicator of platelet activation^28^. We found increased PF-4 concentrations in plasma samples of *Ank^ank/ank^* and especially *Abcc6^−/−^*;*Ank^ank/ank^*mice, suggestive of platelet activation during or after blood collection, which might have obscured our analyses.

**Figure 1:**
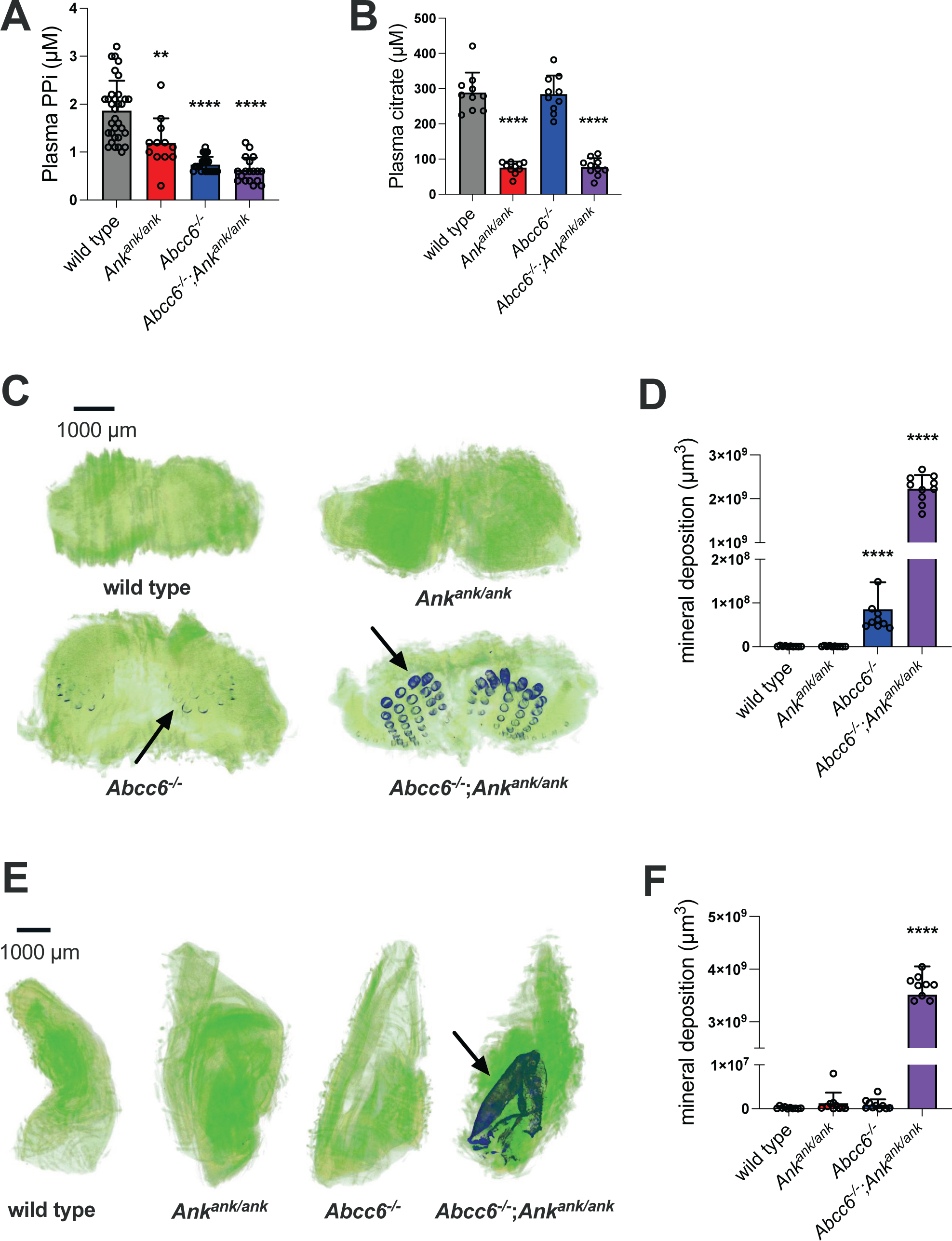
Plasma PPi is regulated by both ABCC6 and ANK, plasma citrate depends solely on ANK, and together these proteins help prevent soft connective tissue calcification. Plasma pyrophosphate (PPi) levels are reduced in *Ank^ank/ank^*, *Abcc6^−/−^*, and double mutant (*Abcc6^−/−^*;*Ank^ank/ank^*) mice compared to wild-type, with the double mutants showing PPi levels similar to *Abcc6^−/−^* mice. (**A**). Plasma citrate concentration is reduced in *Ank^ank/ank^*, and *Abcc6^−/−^*;*Ank^ank/ank^* mice, with levels in *Abcc6^−/−^* mice being not significantly different from wild type. (**B**). Representative microCT images of muzzle skin show calcification of vibrissae capsules (arrows) in *Abcc6^−/−^* mice, which is dramatically increased in *Abcc6^−/−^*;*Ank^ank/ank^*mice. No calcification is detected in wild type or *Ank^ank/ank^*animals (**C**). Quantification of muzzle skin calcification confirms significant increases in ectopic calcification in *Abcc6^−/−^*mice with dramatic ectopic calcification in muzzles of *Abcc6^−/−^*;*Ank^ank/ank^*mice. (**D**) Representative microCT images reveal extensive ectopic calcification of the hyaline cartilage of *Abcc6^−/−^*;*Ank^ank/ank^* ears, with ears of *Abcc6^−/−^*, *Ank^ank/ank^*, and wild type animals being free of ectopic calcification (**E**). Quantification of ear cartilage calcification confirms significant ectopic mineral deposition in ears of *Abcc6^−/−^*;*Ank^ank/ank^* compound mutants, while ears of the other strains remain devoid of calcification. (F). Similar numbers of male and female mice were included in all experiments. Individual data points are shown. Bars represent mean and error bars SD.

### Plasma citrate concentrations depend on ANK

Plasma contains substantial amounts of citrate with reported concentrations in wild type mice ranging from 200-400 µM^25^. Wild type mice in our study had plasma citrate concentrations ranging from 225-421 µM (average 289 ± 56.7 µM) (Fig. 1B). Clearly, most of the citrate in plasma depends on ANK activity, as concentrations in *Ank^ank/ank^* mice were ∼75% lower than in wild type controls (mean in *Ank^ank/ank^* mice 76 ± 17 µM). As expected, plasma citrate levels in *Abcc6^−/−^* mice were indistinguishable from those detected in wild type mice. Similar citrate plasma concentrations were also detected in *Ank^ank/ank^* and *Abcc6^−/−^; Ank^ank/ank^* mice (mean 76 ± 17 µM in *Ank^ank/ank^*mice versus 78 ± 25 µM in *Abcc6^−/−^;Ank^ank/ank^* mice). Collectively, these data indicate that ABCC6 does not contribute to plasma citrate concentrations and are in line with the fact that, different from ANK, ABCC6 has not been implied in cellular release of citrate.

### ABCC6 and ANK are both involved in prevention of soft connective tissue calcification

Calcification of the blood-filled capsule surrounding the whiskers in the muzzle skin is an early biomarker of disease progression in rodent models of ectopic calcification^23^. As expected, microCT imaging did not reveal muzzle skin calcification in wild type mice, whereas robust calcification was seen in muzzle skin of age-matched *Abcc6^−/−^* mice, confirming earlier reports (Fig. 1C, D)^23,29^. Mineral deposits were absent in muzzle skin of *Ank^ank/ank^* mice of the same age which correlated well with the limited reduction in their plasma PPi concentrations. Compared to *Abcc6^−/−^*mice a dramatic increase in mineral deposition was seen in muzzle skin of *Abcc6^−/−^*;*Ank^ank/ank^*mice (mean 8.6 ×10^7^ µm^3^ in *Abcc6^−/−^* mice versus 2.2 ×10^9^ µm^3^ in *Abcc6^−/−^*;*Ank^ank/ank^* mice).

We also assessed calcification of the hyaline cartilage of the ear in the same four mouse strains (Fig. 1E, F). microCT analysis showed extensive calcification in ears of *Abcc6^−/−^*;*Ank^ank/ank^* compound mutant mice, whereas the hyaline cartilage of ears of *Abcc6^−/−^* and *Ank^ank/ank^* single mutant animals was not affected. Collectively, these data demonstrate that both ABCC6 and ANK play crucial and complementary roles in the inhibition of ectopic calcification of soft connective tissues like muzzle skin and ear. As ANK affects both plasma PPi and citrate concentrations, it is currently unclear if the dramatic acceleration of ectopic calcification in *Abcc6^−/−^*;*Ank^ank/ank^*mice is caused by the reduction of extracellular PPi, citrate or a combination.

### Citrate has bioavailability after oral administration and at high doses prevents ectopic calcification in a mouse model of pseudoxanthoma elasticum

Connective tissue calcification is markedly more pronounced in *Abcc6^−/−^;Ank^ank/ank^*than in *Abcc6^−/−^* mice. Citrate chelates calcium and at high concentrations has been shown to prevent calcium phosphate precipitation *in vitro*^30^. We therefore explored if there is a role of this TCA cycle intermediate in inhibition of soft connective tissue calcification. First, we determined the pharmacokinetics of orally administered citrate in mice, using our extensively validated animal model for PXE, the *Abcc6^−/−^* mouse. We found that plasma citrate concentrations were almost doubled 15 minutes after administering an oral dose of 300 mM potassium citrate (10 µl/g body weight), and concentrations remained elevated for several hours (Fig. 2A). These data demonstrated substantial bioavailability of orally administered citrate in mice.

**Figure 2:**
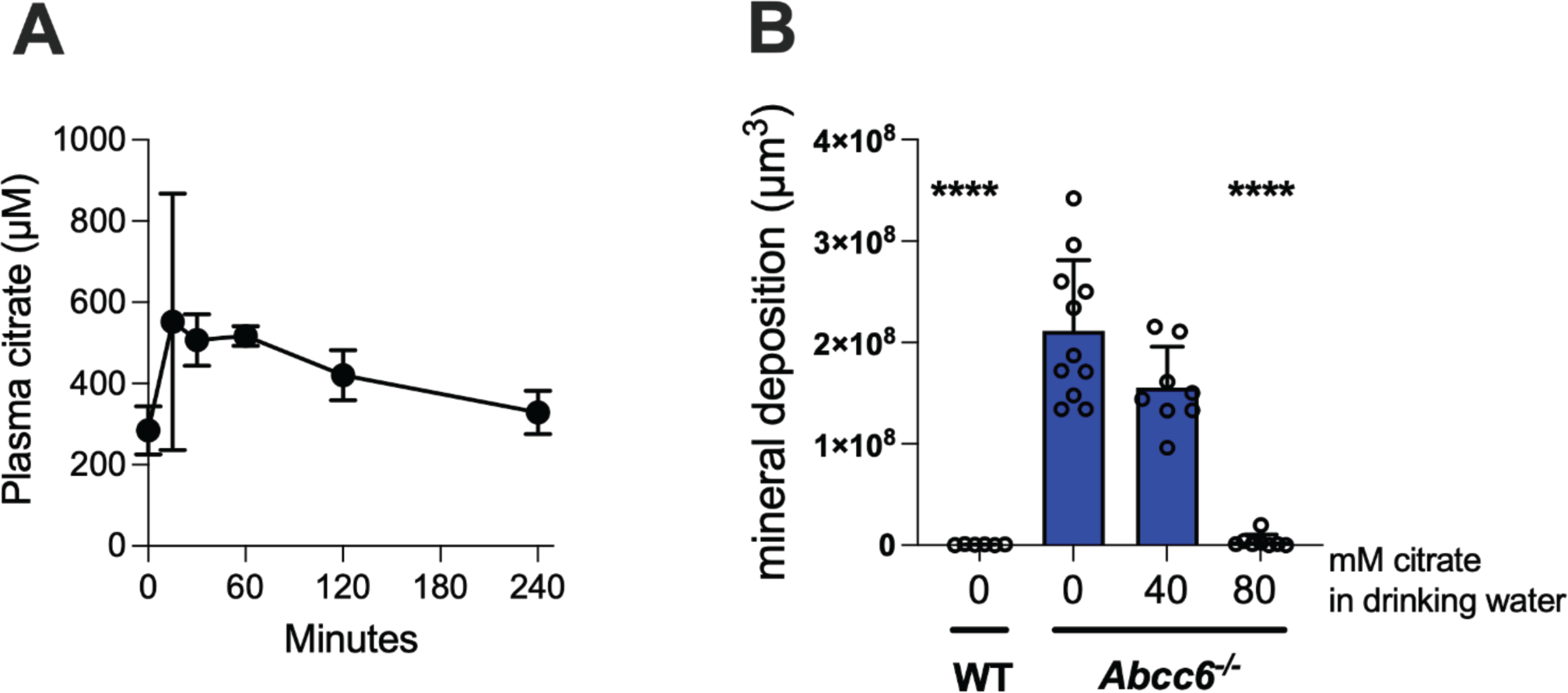
Orally administered citrate is bioavailable and prevents ectopic calcification in *Abcc6^−/−^* mice, a validated model of pseudoxanthoma elasticum. Oral administration of potassium citrate (300 mM, 10 µl/g body weight by gavage) markedly elevated plasma citrate concentrations, which remained increased for several hours, confirming substantial bioavailability of citrate in mice (**A**). Quantification of muzzle skin calcification demonstrates significant, dose-dependent reduction of ectopic calcification in *Abcc6^−/−^* mice upon chronic oral citrate treatment. At the highest dose tested (80 mM), citrate completely prevented ectopic calcification of the muzzle skin (**B**). Similar numbers of male and female mice were included in all experiments. Individual data points are shown. Bars represent mean and error bars SD.

Next, we studied if citrate administered via the drinking water prevented calcification in the muzzle skin of *Abcc6^−/−^* mice. Interestingly, high doses of citrate (80 mM) completely blocked muzzle skin mineralization in *Abcc6^−/−^* mice with a lower dose of 40 mM giving an intermediate effect (Fig. 2B). As orally administered citrate resulted in a significant but not dramatic increase in plasma levels, these data also indicate that endogenous plasma citrate might contribute to inhibition of calcification of soft connective tissues.

### ANK influences both PPi and citrate deposition in bone, while ABCC6 only affects PPi incorporation

The mineral phase of bone contains relatively high amounts of PPi (up to 0.3%, w/w)^31^. We have previously found that ANK-activity underlies about 75% of the PPi incorporated into bone. In our current study, in which we used mice on a C57Bl/6J genetic background, we found a similar reduction in ePPi levels in tibiae of *Ank^ank/ank^* mice (∼ 72% reduction, *Ank^ank/ank^*4.2 ± 0.7 versus wild type 14.5 ± 1.3 nmoles/mg bone) (Fig. 3A). ABCC6 activity was responsible for a smaller amount of the PPi detected in the hydroxyapatite of the tibiae (∼27%) of wild type (*Abcc6^−/−^* 10.6 ± 1.4 nmoles ePPi/mg bone and *Abcc6^−/−^;Ank^ank/ank^* 0.8 ± 0.2 nmoles ePPi/mg bone versus wild type 14.5 ± 1.3 nmoles ePPi/mg bone). Of note, despite their low PPi content, bones of *Ank^ank/ank^* mice do not seem to function as a sink for PPi present in the circulation, as plasma PPi concentrations in *Ank^ank/ank^* mice were only reduced by ∼35%. This is somewhat surprising given the high affinity of PPi for hydroxyapatite and the fact that bone tissue is well perfused.

**Figure 3:**
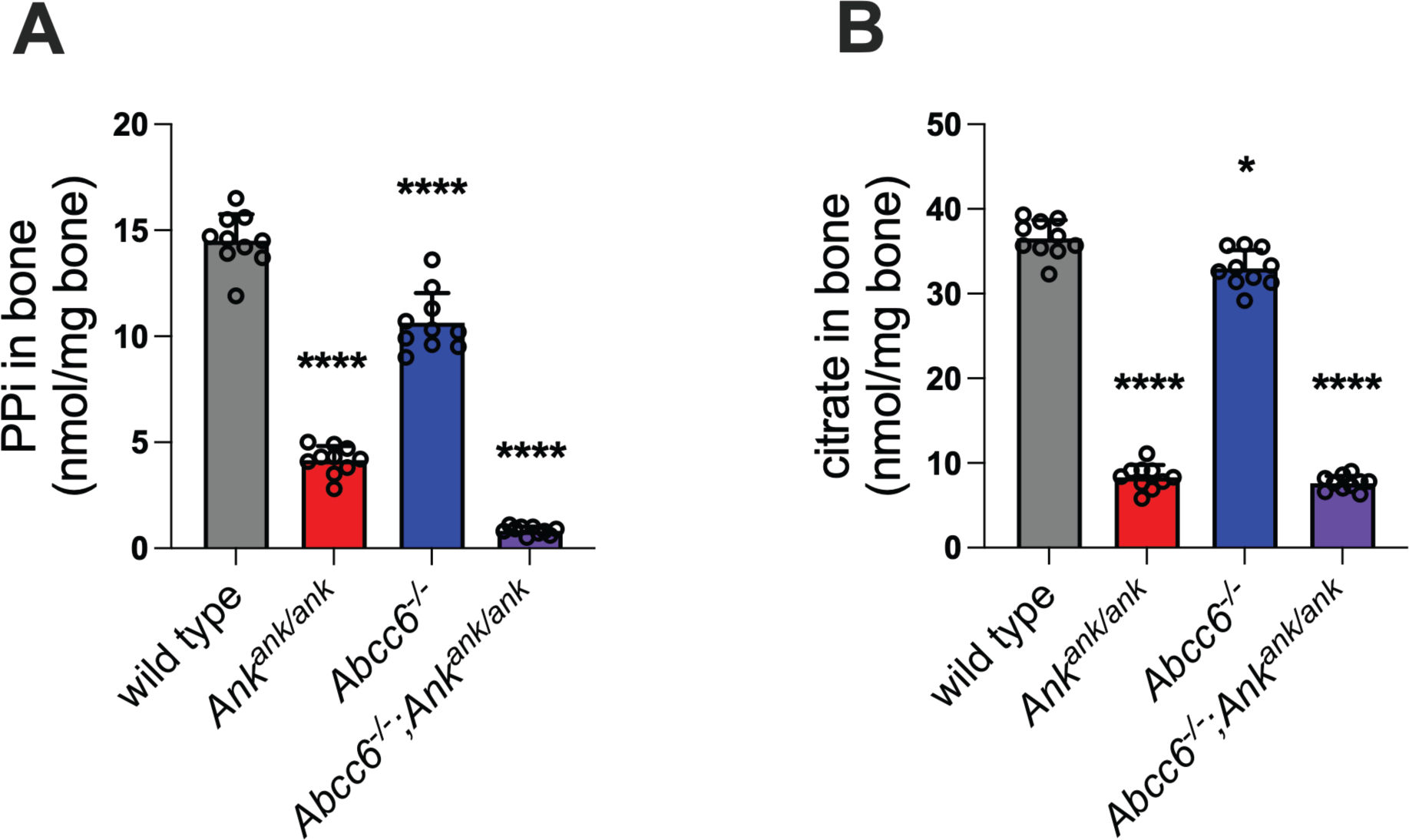
ABCC6 selectively regulates pyrophosphate (PPi), while ANK controls both PPi and citrate levels in bones. Compared to wild type mice, extracellular PPi concentrations in tibiae are markedly reduced in *Ank^ank/ank^* mice with a more moderate reduction seen in *Abcc6^−/−^* mice. In *Abcc6^−/−^;Ank^ank/ank^* compound mutants, tibial PPi is further decreased (**A**). Tibial citrate concentrations are modestly but significantly reduced in *Abcc6^−/−^* mice, whereas *Ank^ank/ank^* and *Abcc6^−/−^;Ank^ank/ank^* compound mutants show comparable and more dramatic reductions (**B**). Individual data points are shown. Bars represent mean and error bars SD.

Citrate is a major organic component of bone, accounting for ∼90% of the body’s total citrate and is strongly associated with hydroxyapatite^32^. We have previously found that ANK-activity underlies ∼ 50% of the citrate incorporated into bone^10^. In the current study, in animals on the C57Bl/6J genetic background, we found an even more robust (∼ 77%) reduction in citrate levels in tibiae of *Ank^ank/ank^* mice (*Ank^ank/ank^* 8.3 ± 1.5 versus wild type 36.6 ± 2.1 nmoles/mg bone) (Fig. 3B). Surprisingly, ABCC6 activity was also responsible for a minuscule but significant amount of the tibial citrate (∼9.7% reduction, *Abcc6^−/−^* 33.0 ± 2.2 nmoles/mg bone versus wild type 36.6 ± 2.1 nmoles/mg bone). The *Abcc6^−/−^;Ank^ank/ank^* showed a significant robust (∼79%) reduction in citrate levels of tibiae (7.6 ± 0.9 nmoles /mg bone versus wild type 36.6 ± 2.1 nmoles/mg bone) similar to that of the single mutant *Ank^ank/ank^* mice, corroborating the strong impact of ANK in determining bone citrate levels.

### Bone structure is influenced by ANK and ABCC6

Reduced levels of extracellular PPi and citrate in *Ank^ank/ank^* and *Abcc6^−/−^*;*Ank^ank/ank^* mice were associated with marked structural alterations in bone, with the most pronounced effects observed in male animals. Loss of ANK led to a significant reduction in trabecular bone volume, as evidenced by a lower BV/TV ratio (*Ank^ank/ank^* 20.3 ± 2.8 and *Abcc6^−/−^;Ank^ank/ank^* 17.0 ± 3.8 % versus wild type 39.8 ± 7.3 %) (Fig. 4). This decrease was driven by both a reduced number of trabeculae and a thinning of individual trabeculae (Supplemental Fig. 1). Alterations in trabecular spacing did contribute much to the lower bone mineral density (BMD) seen in ANK-deficient femora. Intriguingly, compared to *Ank^ank/ank^* mice, trabecular spacing was increased in *Abcc6^−/−^*;*Ank^ank/ank^* mice. The latter effect was seen in male as well as in female animals (Supplemental Fig. 1). In female mice, *Abcc6^−/−^*;*Ank^ank/ank^* animals exhibited a more substantial reduction in trabecular BMD than *Ank^ank/ank^* mice (*Abcc6^−/−^;Ank^ank/ank^* 1.17 ± 0.01 g/cm^3^ and *Ank^ank/ank^* 1.18 ± 0.02 g/cm^3^ versus wild type 1.22 ± 0.02 g/cm^3^). Interestingly, while bones from both genotypes contained similar amounts of citrate, they differed in PPi content (Fig. 3). This raises the possibility that, at least in females, PPi incorporation into bone may play a critical role in regulating trabecular BMD, consistent with the hypothesis proposed by Fleisch and colleagues in the 1970s that PPi stabilizes existing hydroxyapatite deposits^33^. Moreover, these data also point to subtle ABCC6 specific effects on bone structure.

**Figure 4:**
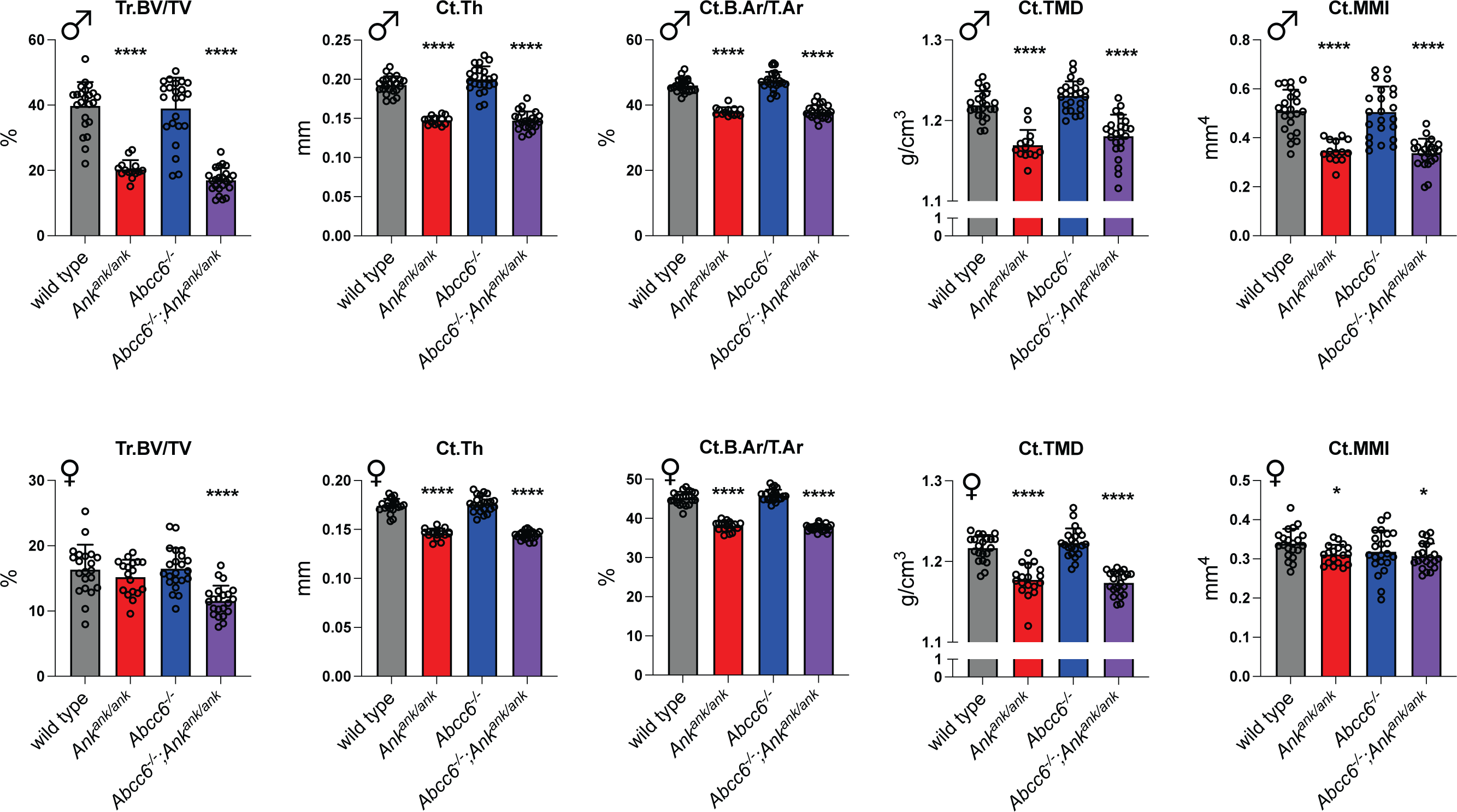
ANK deficiency severely affects bone structure in mice, with effects being more pronounced in females. In male femora, *Ank^ank/ank^* and *Abcc6^−/−^*;*Ank^ank/ank^* genotypes were associated with pronounced structural alterations; a significant reduction in trabecular bone volume and mineralized cortical bone, with decreased cortical thickness and cortical area fraction, next to a reduction in mean moment of inertia. Female ANK-deficient mice, displayed the same, but milder trabecular and cortical bone alterations, across almost all bone structural parameters. Individual data points are shown. Bars represent mean and error bars SD. BMD, Bone Mineral Density; Ct. B.Ar/T.Ar., Cortical Area Fraction; Ct. MMI, Mean Moment of Inertia of Cortical Bone; Ct.Th, Cortical Thickness; Ct. TMD, Cortical Tissue Mineral Density; Tr. BV/TV, Trabecular Bone Volume Fraction

Cortical bone changes in ANK-deficient mice were even more pronounced. These animals showed a substantial reduction in mineralized cortical bone, reflected by decreased cortical thickness (Ct.Th) (*Ank^ank/ank^* 0.148 ± 0.006 mm and *Abcc6^−/−^;Ank^ank/ank^* 0.147 ± 0.012 mm versus wild type 0.193 ± 0.011 mm) and cortical area fraction (Ct.B.Ar/T.Ar) (*Ank^ank/ank^* 38.0 ± 1.3 and *Abcc6^−/−^;Ank^ank/ank^* 37.9 ± 2.1 % versus wild type 46.0 ± 2.1 %). Moreover, cortical bone was severely hypomineralized in the absence of ANK. Notably, these cortical differences were consistent across both male and female mice.

What is more, Mean Moment of Inertia of cortical bone (Ct.MMI) was lower in femora of ANK-deficient animals, indicating altered geometry and reduced resistance to bending forces.

Taken together, our findings highlight a previously underappreciated role for PPi in bone homeostasis, in addition to its well-established function in preventing ectopic calcification of soft connective tissues. We will revisit these seemingly contrasting roles of extracellular PPi in the discussion section.

### ANK and ABCC6 influence the biomechanical properties of long bones

The observed structural differences in long bones across the four mouse strains prompted a detailed analysis of the mechanical properties of their femora. As expected, three-point bending tests revealed significant impairments in bone mechanical performance in animals lacking ANK (Fig. 5). Specifically, Ultimate Moment, Bending Rigidity, and Ultimate Bending Energy were all reduced in femora from ANK-deficient mice. The lower Ultimate Moment (*Ank^ank/ank^* 22.4 ± 1.9 Nmm and *Abcc6^−/−^;Ank^ank/ank^* 23.4 ± 3.0 Nmm versus wild type 36.0 ± 4.6 Nmm in male mice) indicates that these bones are less resistant to fracture compared to those from ANK-proficient animals, which aligns with the observed reductions in cortical thickness and tissue mineral density (TMD). Bending Rigidity, which is strongly influenced by mineralization, was also diminished in *Ank* deficient femora (*Ank^ank/ank^*718 ± 150 Nmm^2^ and *Abcc6^−/−^; Ank^ank/ank^* 698.8 ± 159 Nmm^2^ versus wild type 1150 ± 124 Nmm^2^ in male mice), further corroborating the reduced TMD of cortical bone in these mice.

**Figure 5:**
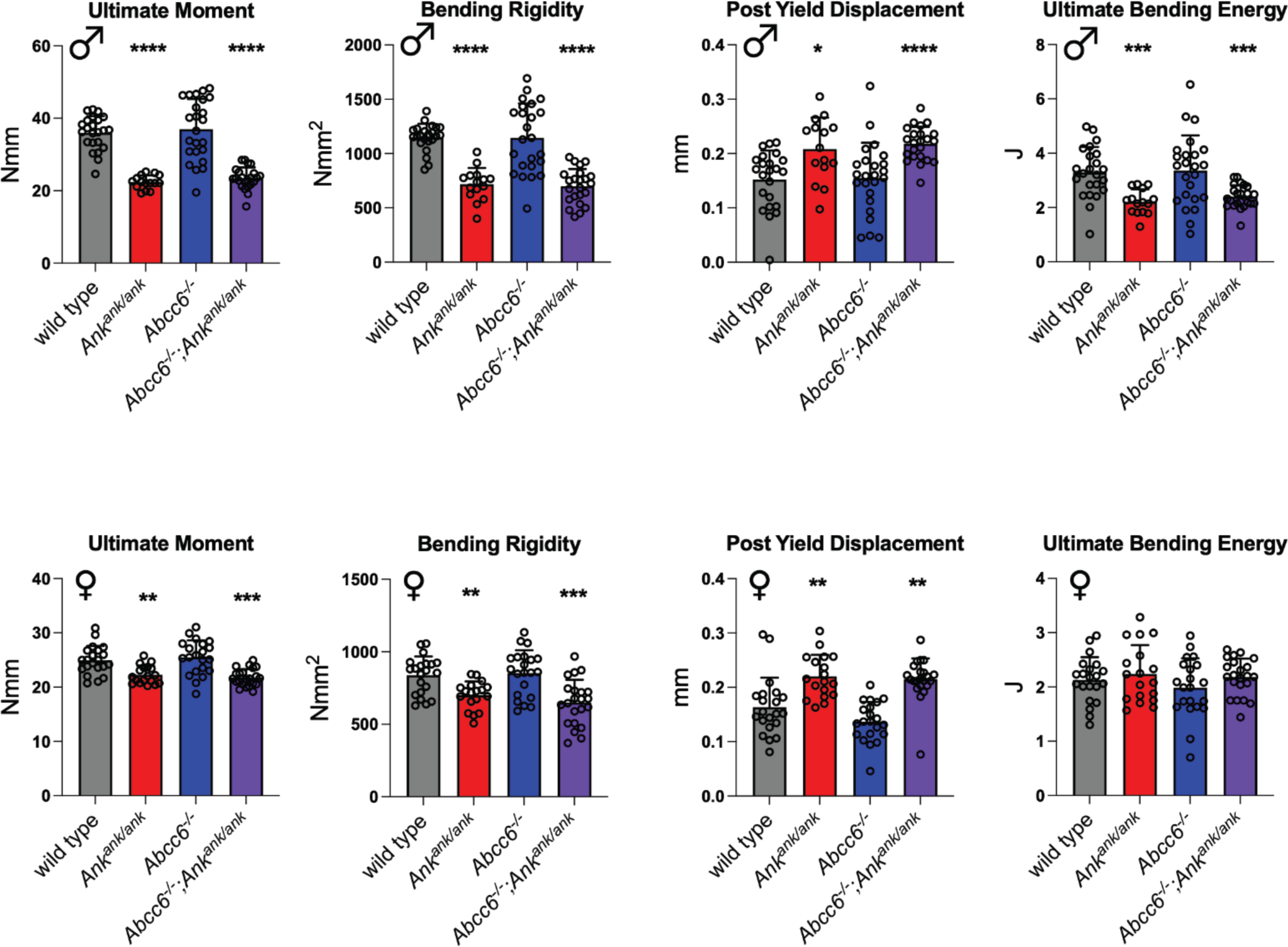
ANK-deficiency severely affects bone biomechanical properties. Three-point bending tests of femora revealed significant impairments in bone mechanical performance in ANK-deficient strains, with effects being most pronounced in males. Ultimate moment and bending rigidity were significantly reduced in *Ank^ank/ank^* and *Abcc6^−/−^;Ank^ank/ank^* mice of both sexes, whereas ultimate bending energy was decreased only in males. In contrast, post-yield displacement was significantly increased *Ank^ank/ank^*and *Abcc6^−/−^*;*Ank^ank/ank^* compound mice of both sexes, indicating compensatory changes in the organic phase of bone. Individual data points are shown. Bars represent mean and error bars SD.

Interestingly, Post-Yield Displacement was increased in both *Ank^ank/ank^*(37% in males and 35% in females) and *Abcc6^−/−^*;*Ank^ank/ank^* mice (31% in males and 35% in females) compared to wild type (0.15 ± 0.05 mm in male and 0.16 ± 0.05 in female), suggesting enhanced strength of the organic matrix in ANK-deficient bones. Notably, in female mice, Ultimate Bending Energy, which is a measure of overall resistance to failure, did not differ significantly across the four strains. In males, differences in this parameter (*Ank^ank/ank^* 2.2 ± 0.5 J, *Abcc6^−/−^* 3.4 ± 1.3 J and *Abcc6^−/−^;Ank^ank/ank^* 2.4 ± 0.4 J versus wild type 3.3 ± 0.9 J in male mice), were less pronounced than those observed for Ultimate Moment. Taken together, these findings suggest that the organic phase in ANK-deficient bones is strengthened, likely as a compensatory response to reduced mineralization.

Mechanical parameters of *Ank^ank/ank^* and *Abcc6^−/−^*;*Ank^ank/ank^*femora were very similar. This indicates that ANK activity is most important for retaining normal bone physiology. The further reduction in PPi incorporation in the mineral phase of *Abcc6^−/−^*;*Ank^ank/ank^* bones apparently does not have a major effect on bone mechanical performance, despite the reduced cortical BMD in female *Abcc6^−/−^*;*Ank^ank/ank^*mice.

## Discussion

Our previous work demonstrated that both ABCC6 and ANK are involved in the cellular release of ATP and therefore play key roles in the generation of extracellular inorganic pyrophosphate (ePPi), a critical regulator of mineralization^10,11,24^. In addition, ANK also mediates the cellular export of citrate thereby affecting citrate levels in plasma, urine, and bone^10^.

Intriguingly, ANK is highly expressed in osteoblasts and osteocytes^12,13^, yet its functional role in bone physiology remains incompletely understood. In this study, we examined the roles of ABCC6 and ANK in preventing ectopic calcification in soft connective tissues and regulating bone physiology. We also explored if citrate, due to its calcium-chelating properties, may contribute to inhibiting ectopic calcification. Notably, we recently found that oral citrate prevents age-related calcification of intervertebral disc structures^34^. Building on this, our current data show that even modest increases in plasma citrate can prevent calcification of muzzle skin in *Abcc6^−/−^* mice. These findings support citrate’s potential as a therapeutic agent and suggest that endogenous citrate may play a natural role in protecting against ectopic calcification.

Plasma PPi levels are regulated by both ABCC6 and ANK, with ABCC6 being the primary contributor^9,11,24^. We observed no significant difference in plasma PPi between *Abcc6^−/−^* and *Abcc6^−/−^*; *Ank^ank/ank^* mice, which we attribute to technical challenges during blood collection. In *Abcc6^−/−^*mice, plasma PPi levels are already low^9,24^, making further reductions difficult to detect. Additionally, ENPP1, which is abundantly present in plasma^35^, rapidly converts any ATP released during blood draw into PPi, potentially skewing measurements. We frequently observed elevated plasma PPi levels due to suboptimal blood collection. Notably, serum PPi concentrations are consistently higher than plasma levels^36^, reflecting ATP release from activated platelets and subsequent conversion to PPi after blood collection. Platelet activation during blood sampling therefore likely affects the concentration of PPi detected in plasma.

Consistent with previous findings, approximately 75% of plasma citrate was found to depend on ANK activity. Although absolute plasma citrate levels were higher in this study, they were measured using NMR, a more robust and quantitative method than the LC/MS approach we used previously^10^. Thus, the current values likely better reflect true plasma citrate concentrations. Citrate, as a divalent cation chelator, may reduce free calcium levels in plasma and thereby inhibit ectopic calcification. Our pharmacokinetic analysis showed that orally administered citrate is absorbed and elevates plasma levels for several hours. Moreover, long-term administration of potassium citrate via drinking water completely prevented ectopic calcification in *Abcc6^−/−^*mice. However, the doses required were high and likely impractical for human PXE patients. Phosphocitrate, a far more potent inhibitor of calcification than either citrate or PPi, may offer a promising alternative^37^. We have initiated studies to evaluate its therapeutic potential, given its significantly higher efficacy in preventing ectopic calcification.

Both ABCC6 and ANK are essential for preventing ectopic calcification in soft tissues. This is evidenced by the significantly greater mineral deposition in the muzzle skin of *Abcc6^−/−^*;*Ank^ank/ank^* mice compared to *Abcc6^−/−^*mice. Similarly, mineralization of the hyaline cartilage in the ear occurred only in mice deficient in both, ABCC6 and ANK, with ears of *Abcc6^−/−^* and *Ank^ank/ank^* mice being indistinguishable from those of wild-type controls. The involvement of ABCC6 in preventing cartilage calcification was unexpected, given the avascular nature of cartilage and ABCC6’s role in regulating PPi in the central circulation. However, due to the thinness of ear cartilage, it is plausible that PPi diffuses from surrounding vasculature into the tissue. While we cannot entirely rule out low-level ABCC6 expression in ear chondrocytes, the low cell density of hyaline cartilage argues against a significant local contribution. The protective effect of ABCC6 against calcification in ear cartilage underscores the ability of circulating PPi to diffuse into peripheral tissues and inhibit ectopic calcification.

Our current data clearly show that ANK plays a protective role against soft tissue calcification. Stimulating ANK activity may therefore offer therapeutic potential for patients with ectopic calcification disorders. Notably, *Abcc6^−/−^*mice with a single inactivated *ank* allele exhibit already dramatically increased calcification over *Abcc6^−/−^* mice with two functional *ank* alleles (manuscript in preparation), further supporting our hypothesis that differences in ANK activity affect ectopic calcification of soft connective tissues. Future studies will focus on identifying small molecules that enhance ANK function, to increase local ATP release and PPi production with aim to halt disease progression in conditions like PXE, which results from ABCC6 deficiency.

PXE patients show striking variability in disease progression, even among individuals with identical ABCC6 mutations within the same family^38,39^. We propose that differences in ANK activity may contribute to this observed phenotypic variability. For instance, environmental factors such as diet and resulting circulating metabolites may influence ANK function. The microbiome may also play a role, as recent work by Mastrangelo et al.^40^ showed that imidazole propionate, a microbial metabolite, promotes atherosclerosis in both humans and mice. It is conceivable that other microbial metabolites modulate ectopic calcification, potentially through interactions with ANK. Interestingly, a recent study found that the gut microbiome of PXE patients differs from that of unaffected individuals^41^. Moreover, patients with more severe phenotypes exhibit reduced bacterial diversity compared to those with milder disease.

ANK is abundantly expressed in osteoblasts and osteocytes^12,13^, and its activity accounts for the majority of PPi incorporated into the mineral phase of bone. Our findings also reveal that ABCC6 contributes to PPi accumulation in bone, likely through plasma-derived PPi that binds tightly to hydroxyapatite. Supporting this notion, is our recent finding that PPi administered via drinking water increases PPi content of long bones, suggesting that circulating PPi can be incorporated into bone tissue^42^. Bone PPi levels are relatively high, comprising approximately 0.1–0.3% of total bone mass^11,31^. At first glance, the presence of a calcification inhibitor like PPi in mineralized bone may seem paradoxical. However, the active incorporation of PPi into bone by ANK-expressing cells suggests a functional role beyond inhibition of ectopic calcification. We hypothesize that PPi contributes to stabilization of existing hydroxyapatite crystals, rather than only simply preventing new mineral formation. This idea is supported by earlier *in vitro* work from the Fleisch group, which demonstrated that PPi-coated hydroxyapatite dissolves more slowly than uncoated crystals^43^. The reduced mineral density observed in bones of ANK-deficient mice aligns well with this hypothesis. Zimmerman et al. ^44^ recently reported that the reduced mineralization of long bones in *Enpp1^asj/asj^* mice, which lack ENPP1 activity, is partly due to the absence of enzymatic activity. The ecto-enzyme ENPP1 primarily generates PPi^45^. Consequently, lower PPi levels in bone tissue are likely to contribute to the diminished mineralization observed in ENPP1-deficient long bones^46^. Intriguingly, Zimmerman and coworkers also found that ENPP1 influences bone mineralization through a mechanism that is independent of its enzymatic activity^44^, though the nature of this non-enzymatic pathway remains unknown.

Although PPi incorporation in long bones was more severely reduced in *Abcc6^−/−^*;*Ank^ank/ank^* than in *Ank^ank/ank^* mice, structural parameters of cortical bone were remarkably similar between the two strains. This suggests that ABCC6 has only a modest impact on bone architecture, and that reducing PPi levels in bone beyond 75% does not exacerbate structural defects. The lack of differences in bone structure between wild-type and *Abcc6^−/−^* mice further supports the conclusion that ABCC6 is not a major regulator of bone morphology, at least in the 3-month-old animals studied here. It is important to note, however, that aged *Abcc6^−/−^* mice have been reported to show reduced mineralization of tibiae^47^ and spinal vertebrae^48^. This may reflect a gradual effect of impaired PPi incorporation into bone over time. While ABCC6’s role in bone is less direct than in soft tissue, its absence could contribute to long-term structural changes.

Mechanical testing via three-point bending revealed that the structural changes in ANK-deficient femora, characterized by reduced mineralization, translated into diminished bone strength. These findings underscore the importance of ANK activity in maintaining normal bone structure and mechanical strength. Our data also suggest that enhancing ANK function may represent a therapeutic strategy for improving bone quality in conditions such as osteoporosis.

The current study established that ANK facilitates the incorporation of substantial amounts of both PPi and citrate into bone mineral. However, the current data do not allow us to distinguish the individual contributions of these two molecules to bone structure. Notably, *Ank^ank/ank^* and *Abcc6^−/−^*;*Ank^ank/ank^* mice accumulate similar levels of citrate in their bones, but their bones differ in PPi content. Despite this, their femoral structural and mechanical properties are nearly identical, which suggests that citrate may play a significant role in regulating bone architecture. However, previous studies complicate this interpretation. For example, ENPP1-deficient mice, which are expected to have normal extracellular citrate homeostasis, exhibit bone defects similar to those seen in ANK-deficient animals^49^. This implies that PPi deficiency alone may be sufficient to impair bone structure. To further dissect the relative contributions of PPi and citrate, we have initiated studies using ENPP1-deficient mice and compound mutants lacking both ENPP1 and ANK (*Enpp1^asj/asj^*;*Ank^ank/ank^*). These models will help clarify the distinct and overlapping roles of these two mineralization regulators in bone physiology and ectopic calcification.

## Acknowledgements

This research was supported by the National Institutes of Arthritis and Musculoskeletal and Skin Diseases under award nos. R01AR082460 and R21AR083597 to KW. Further funding for this work was provided by PXE International to KW. FS received funding of the National Research Development and Innovation Office of Hungary, grant FK131946. The content is solely the responsibility of the authors and does not represent the views of the funding agency.

## Abbreviations

ABCC6: ATP Binding Cassette Subfamily C Member 6
ANK: Progressive Ankylosis Protein
BMD: Bone Mineral Density
Ct. B.Ar/T.Ar.: Cortical Area Fraction
Ct. MMI: Mean Moment of Inertia of Cortical Bone
Ct.Th: Cortical Thickness
Ct. TMD: Cortical Tissue Mineral Density
ENPP1: ectonucleotide pyrophosphatase phosphodiesterase 1
ePPi: extracellular inorganic pyrophosphate
Nmm: Newton-millimeter
N/mm²: Newton per square millimeter
PPi: inorganic pyrophosphate
TCA cycle: Tricarboxylic Acid cycle
TNAP: tissue non-specific alkaline phosphatase
TMD: Tissue Mineral Density
Tr. BV/TV: Trabecular Bone Volume Fraction

---

**Supplemental Figure 1.**
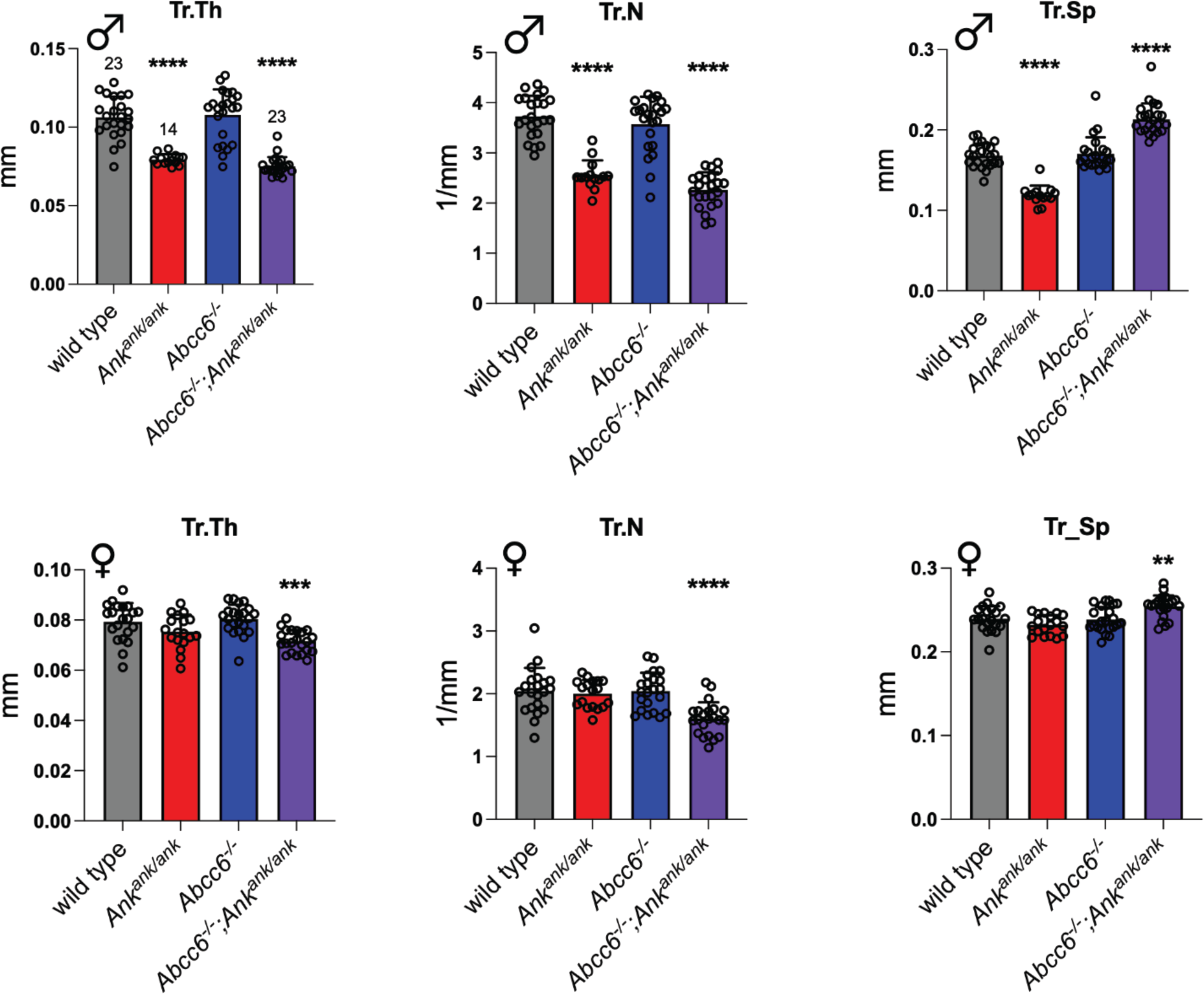

